# Single-section multiplexed imaging enables comprehensive lung cancer diagnosis

**DOI:** 10.64898/2026.04.05.716628

**Authors:** Raz Ben-Uri, Tal Keidar Haran, Yuval Bussi, Gilad Vainer, Johnathan Arnon, Nir Pillar, Stanislav Bahlai, Hasan Sourikh, Inbal Fuchs, Ofer Elhanani, Tzahi Neuman, Eli Pikarsky, Leeat Keren

## Abstract

Accurate and timely diagnosis is essential for effective lung cancer treatment. However, contemporary workflows rely on sequential immunohistochemistry of small biopsy specimens, which can exhaust tissue, limit biomarker assessment, and delay treatment decisions. Here, we demonstrate that multiplexed imaging addresses these limitations by enabling comprehensive lung cancer diagnosis from a single tissue section. We developed and validated a clinically informed multiplexed antibody panel that integrates tumor classification, predictive biomarker assessment, and immune profiling. In diagnostic biopsies, multiplexed imaging achieved 96% concordance with standard pathology, while enabling accurate automated PD-L1 scoring and rapid detection of clinically approved and emerging actionable targets. Simultaneously measuring dozens of proteins improves standard pathology by incorporating complex multi-protein biomarkers, supporting quantitative computational analysis to streamline diagnosis, and generating spatial data for translational research. By reducing turnaround time and preserving scarce tissue, this workflow has the potential to accelerate treatment decisions, improve patient outcomes and bridge clinical care with translational discovery.

## Introduction

Lung cancer is the leading cause of cancer-related mortality worldwide, with approximately 2.5 million new cases and over 1.8 million deaths reported annually ^1^. However, therapeutic advances over the past two decades, including targeted therapies (e.g. EGFR and ALK inhibitors) and immune checkpoint inhibitors (e.g. PD-1/PD-L1 blockade), have markedly improved outcomes for select patient populations ^2–8^. At the same time, refined classification and the growing number of actionable biomarkers have rendered the diagnostic process progressively more complex ^9–11^.

Contemporary lung cancer diagnostics relies on minimally invasive biopsies to reduce patient risk, a shift that has increased dependence on small biopsy specimens. Biopsies are used to establish malignancy, tumor origin, histologic subtype, stage and treatment planning. Because biopsy material is limited, the diagnostic workflow proceeds through a sequential pipeline, beginning with standard hematoxylin and eosin (H&E) staining and immunohistochemistry (IHC) for tumor classification. Histopathological evaluation distinguishes primary lung cancer from other thoracic malignancies or metastatic lesions, separates small cell from non-small cell lung cancer (SCLC/NSCLC), and further subtypes NSCLC into adenocarcinoma, squamous cell carcinoma, and less common variants ^12,13^. These distinctions are essential as therapeutic strategies differ across subtypes ^14^. Subsequently, tissues are tested for predictive biomarkers through additional IHC stains, and next-generation sequencing (NGS) ^15,16^.

While well established, this stepwise approach is inherently slow and constrained by material availability. In complex cases, serial staining may deplete the limited material before essential biomarkers can be evaluated ^12^. Approximately one in five biopsies is ultimately insufficient for a complete molecular workup ^14,17^, necessitating repeat invasive biopsies that increase patient risk and delay therapy initiation. These delays are particularly consequential given that most lung cancer patients present with advanced disease, where early mortality remains high, and rapid and well-informed treatment decisions are essential for survival. Up to 25% of patients die within 1-3 months of diagnosis, often before a full diagnostic evaluation is completed or treatment is initiated ^18–23^. Accordingly, implementing faster diagnostic workflows in this clinical setup can be lifesaving (**Fig. 1A**).

**Figure 1:**
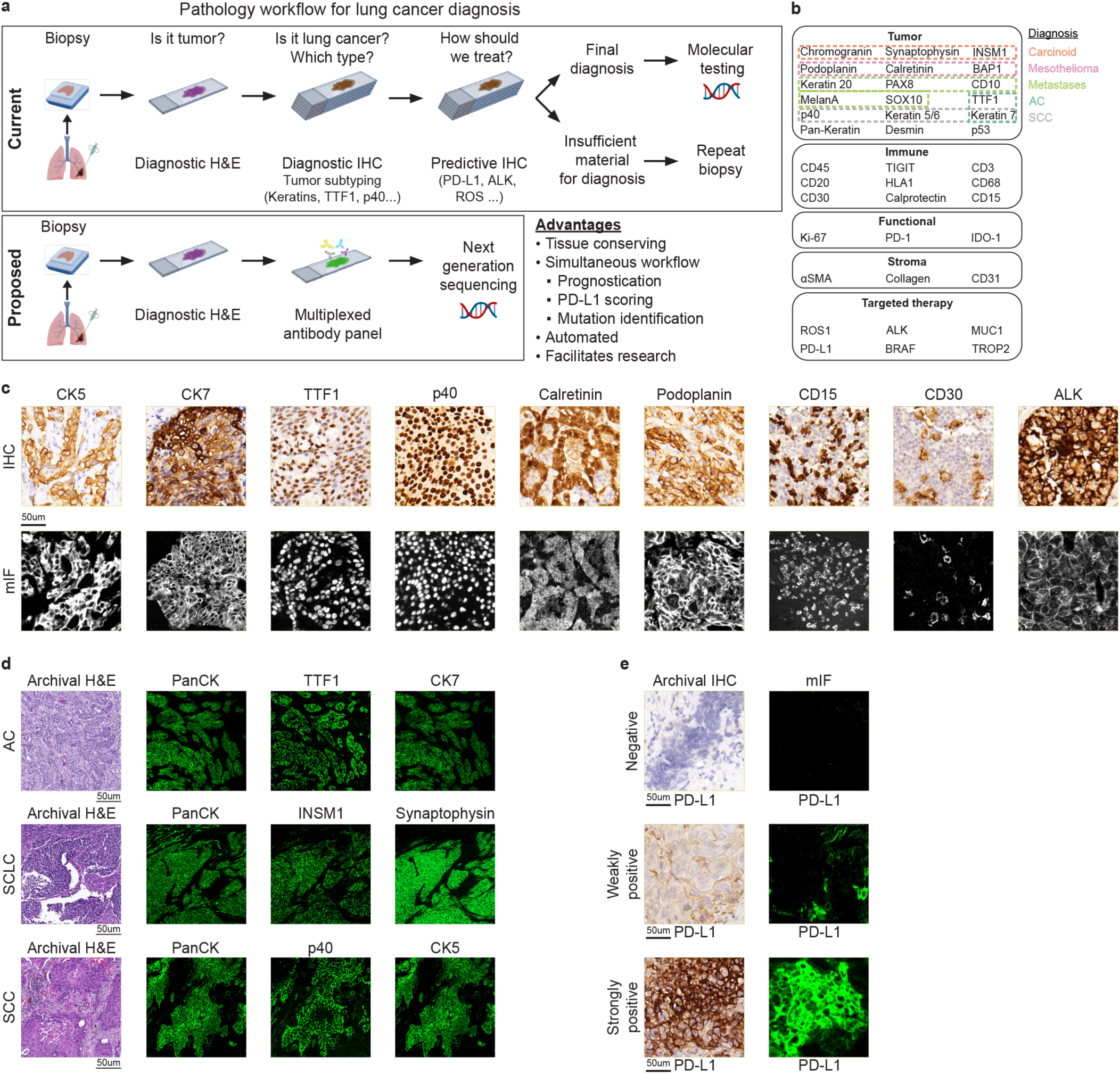
Calibration of multiplexed imaging workflow. **(a)** Conventional immunohistochemistry workflow (top) vs. multiplexed imaging-based workflow (bottom). **(b)** 39-plex antibody panel designed for diagnosis, biomarker evaluation, mutation identification, and immune composition analysis. **(c)** Representative IHC stains (top) and mIF stains (bottom) showing antibody calibration. **(d)** Representative cases demonstrating multiple immunofluorescent stains in addition to H&E, leading to a final diagnosis. **(e)** Archival PD-L1 IHC stains (left) and mIF PD-L1 stains (right) demonstrating all three tumor proportion scores. Abbreviations: AC = Adenocarcinoma, SCC = Squamous cell carcinoma, SCLC = Small cell lung carcinoma.

Beyond delays, sequential IHC introduces interpretive variability. Because markers are assessed on separate serial sections, pathologists must mentally integrate information across slides that are not perfectly aligned. This limitation is exemplified in PD-L1 scoring, which is based on the tumor proportion score (TPS) – the percentage of viable tumor cells exhibiting membranous PD-L1 staining. In routine practice, tumor areas are identified on H&E or lineage-marker stains, and PD-L1 positivity is assessed on a separate slide. With clinically relevant PD-L1 thresholds as low as 1% guiding treatment selection, even minor shifts in tissue architecture between sections can alter the estimated fraction of positive tumor cells and influence treatment selection ^15,24,25^. In addition, tissue exhaustion limits downstream molecular testing and research applications, a challenge that has intensified with the adoption of neoadjuvant therapies. Conserving untreated diagnostic biopsy material for additional testing and translational research is increasingly critical.

Several approaches have been proposed to address these limitations of sequential IHC workflows. Serial multiplexed IHC strategies enable staining of a limited number of markers on a single tissue section ^26–28^. However, expanding from one stain per slide to a handful remains incremental, limiting the impact on routine practice. Other emerging strategies include digital pathology methods that infer protein expression from H&E images ^29,30^ and liquid biopsy approaches that assess circulating tumor DNA or cells ^31,32^. Although promising, these technologies currently complement rather than replace tissue-based assays and still require direct protein measurements for diagnostic decision-making. Similarly, while ultra-fast NGS platforms are advancing, broad implementation remains limited^33,34^. Clinical guidelines recommend a target turnaround time of ≤10 days from biopsy to actionable results ^35^. However, real-world studies in NSCLC report median times of approximately 6 days to histologic diagnosis, and additional 2-5 days to PD-L1, and around 3 more weeks for NGS with total time from biopsy to treatment initiation often extending to 35-45 days ^36–41^. Turnaround times at our institution are comparable (Table S1), reflecting persistent workflow constrains inherent to sequential diagnostic and molecular testing.

Taken together, IHC remains indispensable for tumor classification and biomarker assessment ^42–44^, but its sequential nature and cumulative tissue consumption underscore the need for more efficient diagnostic strategies that support biomarker complexity, maintain spatial integrity, minimize tissue exhaustion and accelerate clinical decision-making and discovery. Novel multiplexed imaging technologies offer a potential solution by enabling simultaneous detection of dozens of markers within a single tissue section^45–49^. While widely adopted in research settings, particularly for immune profiling^50–52^, their diagnostic accuracy and clinical utility have not been systematically evaluated.

Here, we demonstrate the application of multiplexed imaging for comprehensive lung cancer diagnosis. We developed a clinically informed antibody panel incorporating markers routinely used for lung tumor subtyping and predictive biomarker evaluation. We performed extensive validations of this panel on diverse lung resections, including multiple NSCLC subtypes, SCLC, metastatic lesions and other thoracic malignancies. We subsequently performed a blind evaluation on biopsy specimens, where it achieved 96% diagnostic concordance with conventional IHC. We leveraged the high-dimensional nature of the data to perform automated PD-L1 quantification. In a blinded assay, multiplexed imaging achieved 97% concordance with PD-L1 tumor proportion scores (TPS) reported by an accredited clinical pathology laboratory, while expediting and standardizing the analysis. Integration of actionable and investigational biomarkers, together with tumor microenvironment (TME) profiling, illustrated how multiplexed imaging can consolidate diagnosis, biomarker assessment, and translational insight within a single assay, seamlessly bridging clinical care with discovery. Collectively, these findings support multiplexed imaging as a tissue-efficient, accurate, and scalable framework for next-generation lung cancer diagnostics.

## Results

### Design and optimization of a multiplexed antibody panel for lung cancer diagnosis

We developed a 39-plex antibody panel for diagnostic evaluation of a broad spectrum of lung pathologies, designed to mirror the conventional clinical workflow and aligned with the guidelines of the American Society of Clinical Oncology (ASCO) and the College of American Pathologists (CAP). The panel integrates markers required to classify primary lung malignancies, such as adenocarcinoma, squamous cell carcinoma, SCLC, and mesothelioma (e.g., TTF-1, p40, CK7, Synaptophysin, Chromogranin, INSM1), as well as lung metastases, including breast, colorectal and renal cell carcinoma (e.g., CD10, CK20). In addition, the panel included biomarkers relevant for stratification of approved therapies, (e.g. PD-L1, ALK, ROS1), emerging therapeutic targets, (e.g. TROP2, MUC1), and immune markers for translational TME profiling (e.g. CD45, HLA-I, CD3). Together, this design enables simultaneous assessment of tumor origin, histologic subtype, predictive biomarkers and immune context within a single staining assay (**Fig. 1A,B**).

We designed the assay to maximize compatibility with clinical workflows. We implemented it on the COMET platform, a cyclic immunofluorescence (IF) system that supports multiplexed imaging of formalin-fixed paraffin-embedded (FFPE) tissue using off-the-shelf primary antibodies and fluorescent secondary antibodies ^53^. This approach enabled direct use of clinical antibody clones without conjugation, facilitating direct translation from standard IHC.

To calibrate the panel, each antibody was individually titrated and validated on FFPE control tissues using both chromogenic IHC and multiplexed IF to confirm specificity, signal intensity, and appropriate subcellular localization (**Fig. 1C; Supplementary Fig. 1A**). We then applied the panel to lung resection specimens from 55 patients representing the major diagnostic categories encountered in clinical practice, refining staining conditions to ensure consistent performance across histologic subtypes and diverse specimens (**Fig. 1D; Supplementary Fig. 1B,C**). For PD-L1, staining sensitivity and specificity were benchmarked against matched PD-L1 tumor proportion scores (TPS) reported by the accredited clinical pathology laboratory as part of routine standard-of-care testing, to ensure concordance in tumor proportion scoring across clinically relevant thresholds (**Fig. 1E**). Collectively, these steps established a clinically compatible, analytically validated 39-plex panel encompassing diagnostic, predictive, and translational markers for lung cancer.

### Multiplexed imaging enables accurate classification of diagnostic biopsies

We tested the performance of the multiplexed assay in an independent cohort of diagnostic biopsies. The cohort comprised 54 transbronchial, endobronchial and core needle biopsies obtained from patients who underwent clinical workup for suspected thoracic malignancy. The cohort encompassed the spectrum of pathologies encountered in clinical practice, including primary lung malignancies (adenocarcinoma, squamous cell carcinoma, small cell carcinoma, carcinoid tumors, and mesothelioma), metastatic tumors (melanoma, renal cell carcinoma, and sarcomas), as well as non-neoplastic and inflammatory conditions **(Fig. 2A)**.

**Figure 2:**
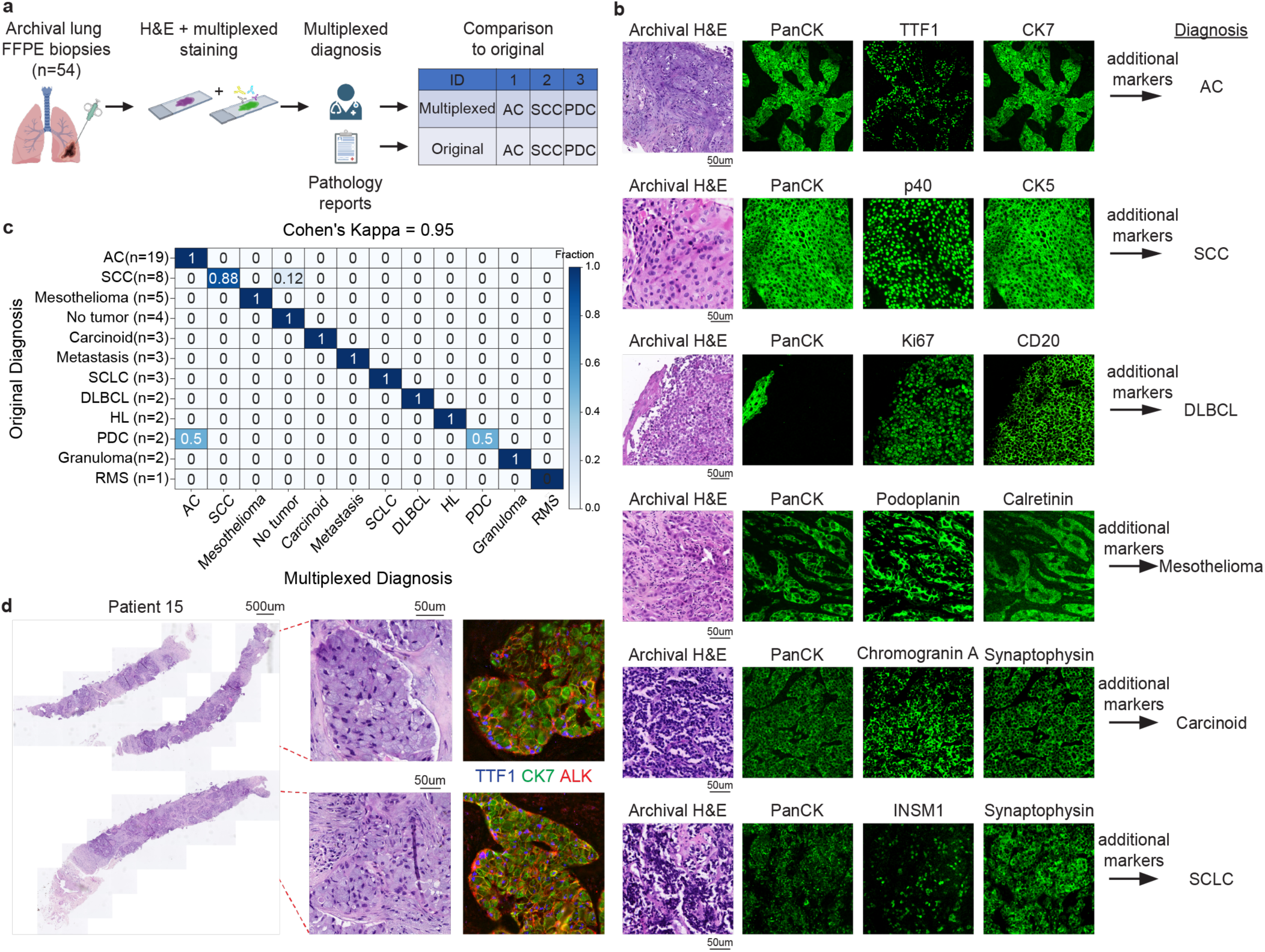
Multiplexed imaging enables accurate classification of biopsies. **(a)** Lung FFPE biopsies were stained for H&E and multiplexed imaging and diagnosed. Results were compared to the original diagnoses. **(b)** Representative cases demonstrating multiple immunofluorescent stains in addition to H&E, leading to a final diagnosis. **(b)** Agreement between multiplexed-based diagnosis (x-axis) and the original diagnosis (y-axis). Color depicts fraction of cases per row. **(d)** Diagnosed poorly differentiated carcinoma showing ALK expression in the multiplexed image. Abbreviations: AC = Adenocarcinoma, SCC = Squamous cell carcinoma, DLBCL = Diffuse large B-cell lymphoma, SCLC = Small cell lung carcinoma, HL = Hodgkin lymphoma, RMS = Rhabdomyosarcoma, PDC = Poorly differentiated carcinoma.

A single section from each biopsy was profiled using the multiplexed protocol. A board-certified pathologist, blinded to the original diagnosis and all clinical metadata, reviewed the multiplexed images alongside paired H&E slides and rendered a diagnosis based on marker expression and morphology. The process from staining to diagnosis took four days. Multiplexed-based diagnoses were then compared with the original clinical diagnoses from the medical records established using conventional H&E and IHC (**Fig. 2A**). Using a single tissue section per case, and blinded to the patients’ clinical data, multiplexed imaging achieved 96% concordance (52/54 cases, Cohen’s κ = 0.95), including accurate classification of rare tumor subtypes and metastatic lesions (**Fig. 2B-C, Supplementary Fig. 2A**).

We identified two discordant cases. In one squamous cell carcinoma case, no tumor was present in the section taken for multiplexed imaging, leading to classification as “no tumor.” Review of adjacent clinical sections confirmed that the tumor area had been exhausted during prior routine diagnostic sectioning of the archived block. This finding highlights the susceptibility of sequential slide-based workflows to tissue depletion and underscores the advantage of staining for many markers on a single section to preserve limited material.

In the second case, the original diagnosis classified the tumor as a poorly differentiated carcinoma based on weak TTF-1 staining. While multiplexed imaging confirmed weak TTF-1 expression, it also revealed strong ALK expression, an actionable alteration that is predominantly associated with adenocarcinoma. In the context of morphology and marker profile, this finding supported reclassification of the tumor as adenocarcinoma (**Fig. 2D**). This case demonstrates how simultaneous assessment of tumor morphology, lineage markers, and predictive biomarkers within a single assay can refine classification in diagnostically ambiguous biopsies. Collectively, these findings demonstrate that multiplexed imaging can accurately classify lung cancer biopsies using a single tissue section, while preserving material for downstream analyses.

### Multiplexed imaging enables accurate manual and automated PD-L1 scoring on a single tissue section

PD-L1 expression is a key biomarker guiding immunotherapy treatment in lung cancer and is routinely assessed by IHC using the tumor proportion score (TPS), defined as the percentage of viable tumor cells exhibiting membranous PD-L1 staining. TPS stratifies tumors into clinically relevant categories, including negative (<1%), weakly positive (1–49%), and strongly positive (≥50%). Because multiplexed imaging assesses PD-L1 on the same section used to identify tumor and immune cells, we evaluated whether our approach could reproduce TPS scoring and facilitate automated analysis.

To benchmark multiplexed imaging against clinical standards, a board-certified pathologist blindly analyzed 31 diagnostic biopsy specimens spanning all three TPS categories. The pathologist performed manual TPS scoring on the multiplexed images by estimating the fraction of PD-L1–positive tumor cells, while excluding PD-L1–positive immune cells using lineage-specific markers such as CD45 and CD68. The same thresholds applied in the clinic were used to define negative (<1%), weakly positive (1–49%), and strongly positive (≥50%) samples (**Fig. 3A**).

**Figure 3:**
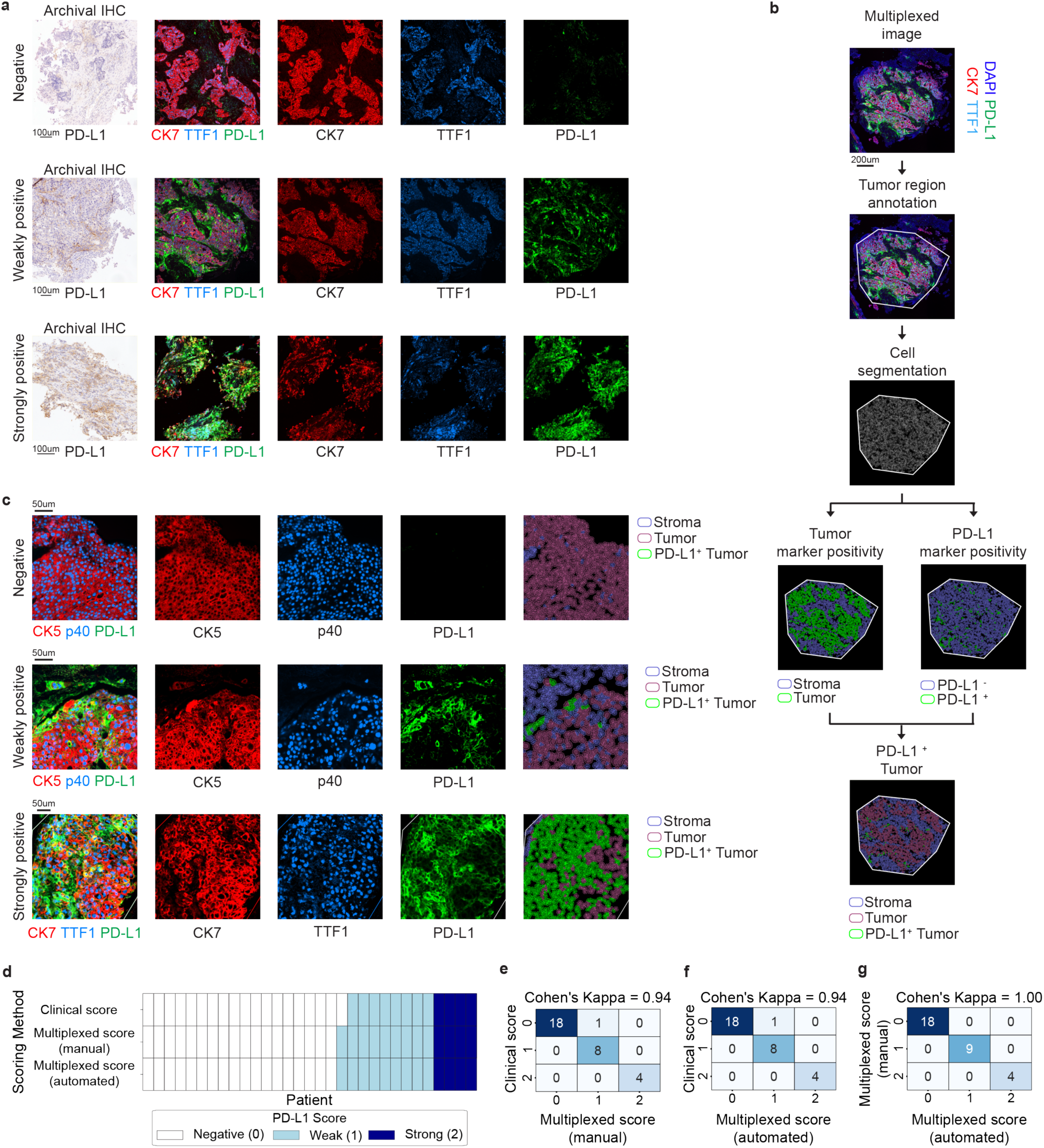
Multiplexed imaging enables accurate manual and automated PD-L1 scoring. **(a)** Representative cases demonstrating manual PD-L1 TPS scoring of negative (<1%, top), weakly positive (1-49%, middle), and strongly positive (≥50%, bottom) tumors, in addition to the original PD-L1 IHC stains. **(b)** Workflow for automated PD-L1 scoring using multiplexed images and marker positivity. A pathologist annotates the tumor regions, followed by cell segmentation based on nuclear staining. Next, tumor and PD-L1 marker positivity are determined using a threshold classifier. Finally, PD-L1-positive tumor cells are determined as the intersection between the two classifiers. **(c)** Representative cases demonstrating automated PD-L1 TPS scoring of negative (<1%, top), weakly positive (1-49%, middle), and strongly positive (≥50%, bottom) tumors. Stroma (blue), PD-L1 negative tumor (light red), PD-L1 positive cells (green). **(d)** PD-L1 scoring method (y-axis) tile map by patients (x-axis). PD-L1 scores: 0 = Negative, 1 = Weakly positive, 2 = Strongly positive. **(e)** Confusion between the original PD-L1 score (y-axis) and manual multiplexed score (x-axis). Color indicates number of cases. **(f)** Confusion between the original PD-L1 score (y-axis) and automated multiplexed score (x-axis). Color indicates number of cases. **(g)** Confusion between the manual multiplexed PD-L1 score (y-axis) and automated multiplexed score (x-axis). Color indicates number of cases.

To further enhance reproducibility and reduce subjectivity, we developed an automated TPS scoring pipeline based on the multiplexed data. Tumor regions were annotated to exclude tissue edges and necrotic regions, followed by single-cell segmentation using InstanSeg ^54^. Two independent classifier thresholds were applied: one to identify tumor cells based on tumor-specific markers such as CK7, TTF-1, CK5 and p40, and a second to classify PD-L1 positivity at the cellular level. The intersection of these populations yielded an automated TPS value for each sample (**Fig. 3B, Methods**). We applied this pipeline to the same 31 samples used for manual scoring (**Fig. 3C**).

Comparison of multiplexed-based scoring with the original clinical TPS reports demonstrated high concordance. Both manual and automated multiplexed-based TPS using the same thresholds matched the clinical classification from the patients’ medical records in 30 of 31 cases (97% concordance; Cohen’s κ = 0.94; **Fig. 3D–G**). Manual and automated multiplexed-based scores were fully concordant across all samples (31/31 cases; Cohen’s κ = 1.0; **Fig. 3F**). One sample was discordant between the multiplexed and original clinical assessment (**Fig. 3D)**. In this case, the original TPS classified the tumor as negative for PD-L1, whereas quantitative multiplexed analysis yielded a TPS of approximately 1.5%, placing the tumor just above the clinical threshold for weak positivity (>1%). Review of the medical records indicated that the patient was initially treated with chemoradiation, subsequently developed recurrent disease, and then received nivolumab (anti PD-1) with a favorable response. Together, these results demonstrate that multiplexed imaging supports accurate and reproducible PD-L1 scoring on a single tissue section, enables objective automated TPS quantification, and facilitates resolution of challenging cases near clinically relevant decision thresholds.

### Multiplexed imaging enables detection of clinically actionable biomarkers

We next evaluated whether the multiplexed panel could detect clinically actionable biomarkers relevant for treatment stratification. The panel included antibodies targeting established alterations used in routine clinical practice, such as ALK and ROS1, as well as additional biomarkers of emerging therapeutic relevance, including TROP2, MUC1 and p53 ^55–58^. In current workflows, these biomarkers are evaluated using a combination of IHC and molecular assays such as fluorescence *in situ* hybridization (FISH) or NGS. While NGS provides definitive genomic characterization, it typically requires multiple unstained tissue sections and extended turnaround times. This is particularly consequential for alterations such as ALK and ROS1 rearrangements, which are often associated with aggressive disease courses yet predict high response rates to targeted therapies ^59–61^. In contrast, IHC detects surrogate changes in protein expression, but is faster. We therefore evaluated whether simultaneous assessment of these targets within the multiplexed assay could provide actionable protein-level information earlier in the diagnostic process.

To assess the accuracy of multiplexed biomarker detection, we compared protein expression in the multiplexed images with available clinical NGS results. All three clinically confirmed ALK-positive tumors were correctly identified as over-expressed by the multiplexed assay (100% concordance, **Fig. 4A-B**). Similarly, all 18 tumors with available p53 or ROS1 status were correctly classified (100% concordance, **Supplementary Fig. 2B-E**), showing protein expression patterns consistent with their known genomic alterations. In addition, multiplexed imaging revealed high expression of TROP2 in 10 tumors and MUC1 in 23 tumors within this cohort (**Fig. 4C-D**). Although these biomarkers are not routinely assessed in standard diagnostic workflows, they are under active clinical investigation as targets for antibody-drug conjugates, including recent regulatory approval of a TROP2-directed therapy for refractory EGFR-mutated NSCLC^62,63^, and may serve as biomarkers for ongoing clinical trials^64,65^.

**Figure 4:**
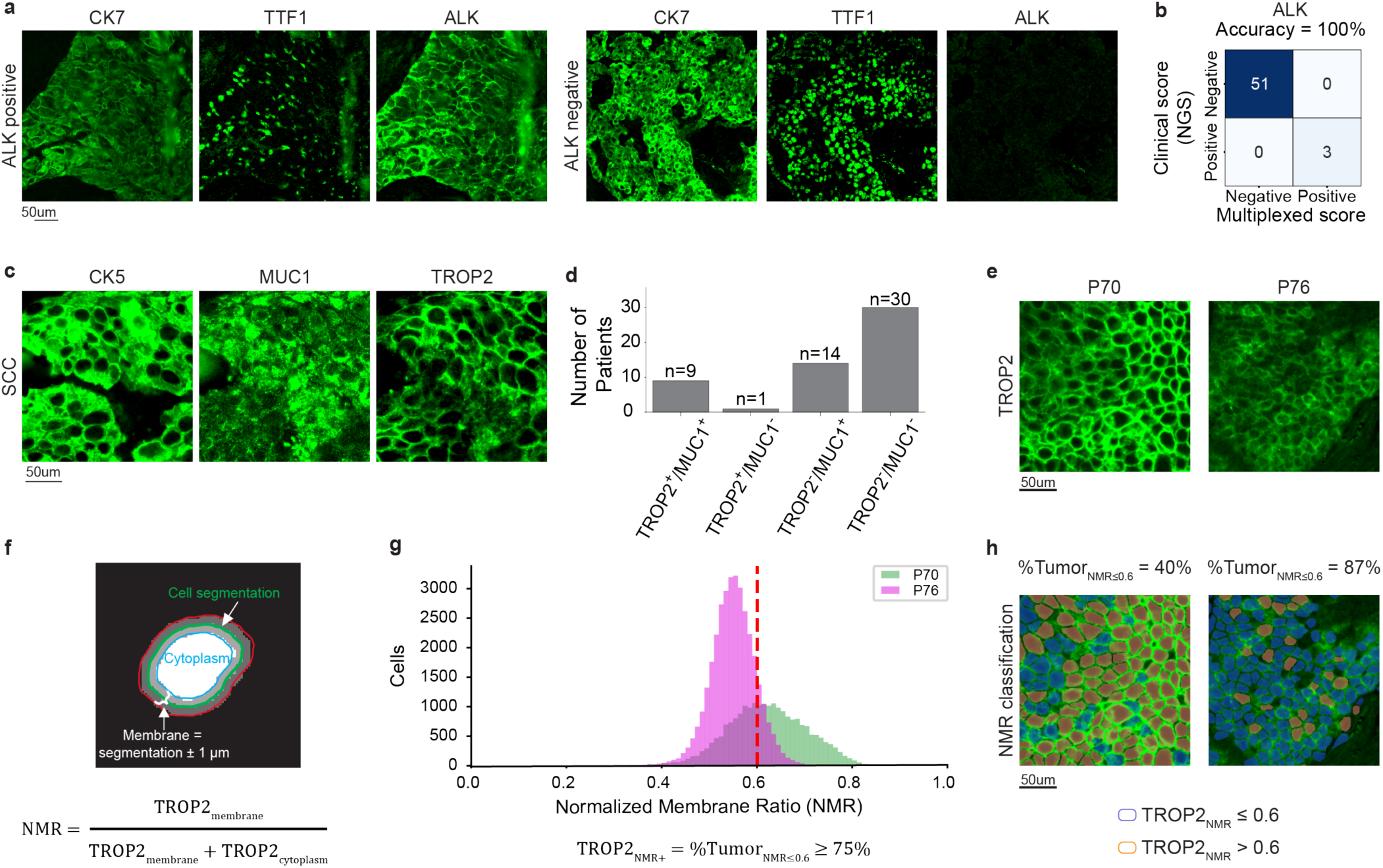
Multiplexed imaging enables detection of clinically actionable biomarkers and mutations. **(a)** Representative images of ALK positive (left) and negative (right) tumors. **(b)** Confusion between NGS (y-axis) and multiplexed imaging (x-axis) in detecting ALK mutation. Color indicates number of cases. **(c)** Representative image of TROP2^+^/MUC1^+^ squamous cell carcinoma. **(d)** Bar graph of TROP2 and MUC1 expression (x-axis) across patients (y-axis). **(e)** Representative images of TROP2-positive patients with two distinct expression patterns. **(f)** NMR (Normalized Membrane Ratio) calculation for TROP2 expression was done using TROP2 membrane measurement (±1µm cell segmentation) and TROP2 cytoplasm measurement. **(g)** Cells count (y-axis) NMR distribution (x-axis) of patients 70 (green) and 76 (magenta). TROP2-NMR+ was defined as tumors with ≥ 75% of cell with TROP2-NMR ≤0.6. **(h)** Same as e, but with NMR classification. Blue = TROP2-NMR ≤ 0.6, red = TROP2-NMR > 0.6.

Multiplexed imaging enables evaluation of subcellular protein localization, which is relevant for biomarkers whose therapeutic efficacy depends on target distribution. For TROP2, low membrane-associated expression quantified by the normalized membrane ratio (NMR) has been shown to predict response to TROP2-directed antibody–drug conjugates (ADCs), and recently approved as a companion diagnostic^66^. To evaluate TROP2 localization, we quantified membrane and cytoplasmic signal intensities from multiplexed images and computed an NMR, defined as the fraction of TROP2 signal localized to the membrane (**Fig. 4E-F, Methods**). This metric is conceptually aligned with the previously-described QCS-NMR score for IHC-based TROP2 evaluation^67^, but is implemented here using immunofluorescence rather than chromogenic staining. We found that a threshold of 0.6 (closely matching the published QCS-NMR cutoff of 0.56) distinguished membrane-low from membrane-high expression states (**Fig. 4G-H**). Although additional validation is required to establish this approach as a predictive biomarker for TROP2-directed ADCs, these findings suggest that multiplexed imaging can support cell-resolved assessment of target localization.

Together, these results demonstrate that multiplexed imaging enables simultaneous assessment of established and emerging actionable biomarkers within a single tissue section, broadening the treatment-relevant information obtainable from limited biopsy material.

### Multiplexed imaging integrates diagnostics and research to enable quantitative analysis of the tumor microenvironment

Finally, we examined whether multiplexed imaging facilitates integration of routine diagnostics with translational research. To this end we incorporated into the panel antibodies to profile the TME, including markers for stromal components, such as CD31, collagen and aSMA, as well as immune cell populations, including CD3, CD20 and CD68. Using InstanSeg^54^ for cell segmentation and CellTune^68^ for cell classification, we classified cells into 14 major epithelial, stromal, and immune populations across all samples (**Fig. 5A**). This framework enabled generation of a standardized immune-cellular report for each patient, summarizing cell-type abundance and spatial distribution, as well as cohort-level statistics (**Fig. 5B-C**). As an example of quantitative immune profiling, we automatically quantified the fraction of tumor-infiltrating cytotoxic lymphocytes (CD8 TILs) for each patient and observed substantial inter-patient variability (**Fig. 5D–E**). While cytotoxic TILs have been shown to carry prognostic and predictive value in lung cancer beyond PD-L1 TPS^69,70^, they are not routinely assessed in clinical practice due to lack of standardized scoring criteria and prospective validation. Here, multiplexed imaging enabled objective, automated cytotoxic TIL quantification directly from diagnostic specimens.

**Figure 5:**
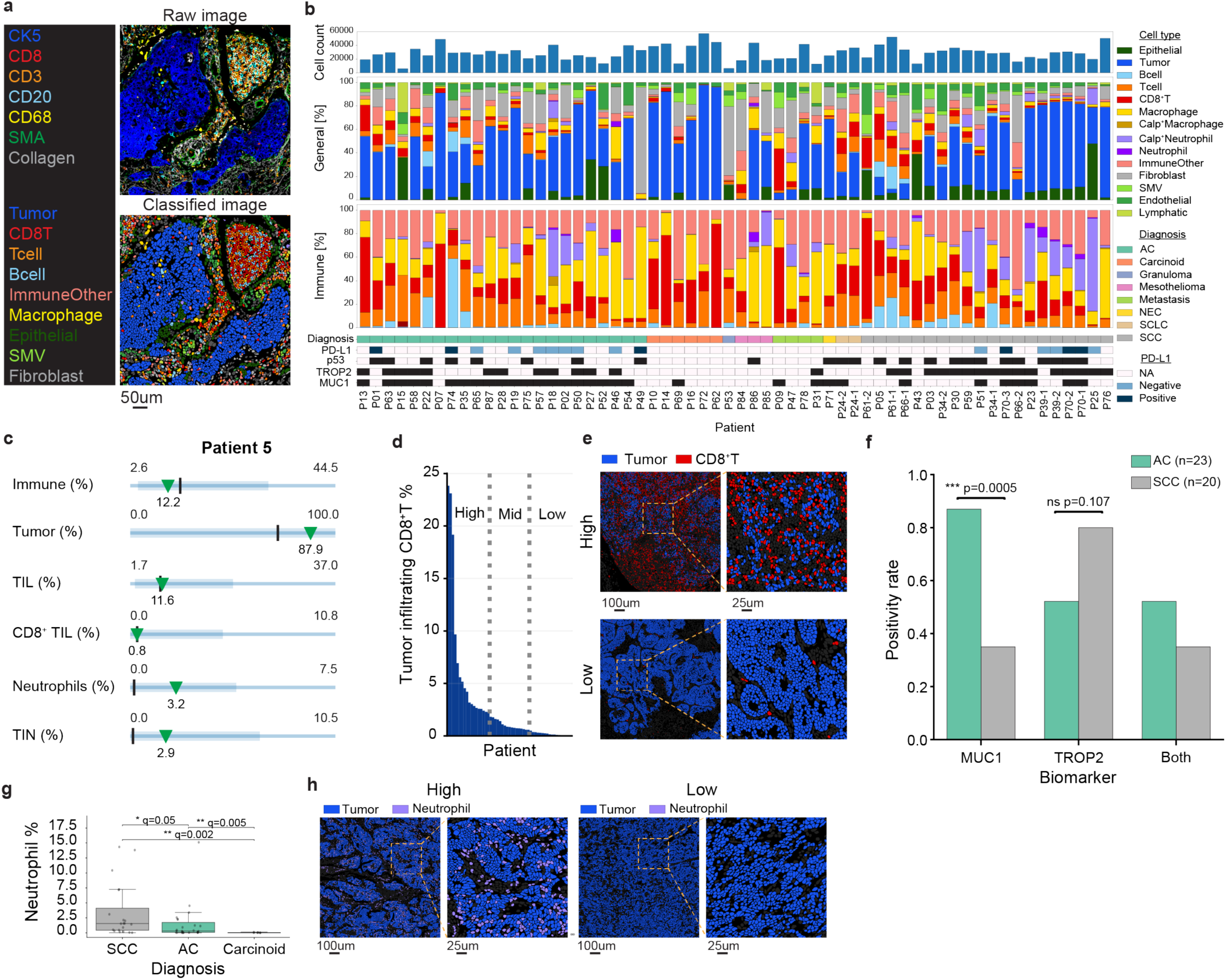
Multiplexed imaging enables quantitative analysis of the tumor microenvironment. **(a)** Protein expression images (top) are used to generate cell-type classifications (bottom) for downstream quantitative analysis. **(b)** Top: total number of cells and their relative abundances (y-axis) across all patients (x-axis) according to tumor subtypes. Middle: the same for immune cells only. Bottom: tumor characteristics per patient. Black is positive for biomarker. **(c)** Example of a standardized cellular report for one patient. Green triangles indicate the patient’s value, black lines indicate the median across patients, blue lines indicate the full range across patients and the thick blue lines indicate 5th-95th percentile. **(d)** Stratification of patients (x-axis) into tertiles according to CD8^+^ T-cell infiltration (y-axis). **(e)** Representative cell classification images of patients with high (top) and low (bottom) CD8^+^ T-cell infiltration. Tumor cells (blue), CD8^+^ T cells (red), other cell types (gray). **(f)** Positivity rate (y-axis) of MUC1 (left), TROP2 (middle), and both (right) in adenocarcinomas (AC) and squamous cell carcinomas (SCC). Statistical significance was assessed using Fisher’s exact test. **(g)** Fraction of neutrophils (y-axis) in squamous cell carcinomas (SCC), adenocarcinomas (AC), and typical carcinoid (TC) tumors (x-axis) with Mann Whitney q values (FDR corrected p values). **(h)** Representative cell classification images of patients with high (top) and low (bottom) neutrophil %. Tumor cells (blue), neutrophils (magenta), other cell types (gray). Abbreviations: AC = Adenocarcinoma, NEC = Neuroendocrine carcinoma, SCLC = Small cell lung carcinoma, SCC = Squamous cell carcinoma.

Beyond cytotoxic TILs, multiplexed profiling revealed histology-associated differences in biomarker expression and immune composition. Consistent with prior reports^71–73^, MUC1 expression was significantly higher in adenocarcinomas compared with squamous cell carcinomas (p = 0.0006), while TROP2 showed a trend toward higher prevalence in squamous cell carcinomas (p = 0.107). Tumors co-expressing both markers did not differ significantly between histologic subtypes (**Fig. 5F**). In addition, squamous cell carcinomas exhibited significantly higher neutrophil fractions than adenocarcinomas (q = 0.05), which in turn were higher than those observed in carcinoid tumors, with a similar result when quantifying the fraction of tumor infiltrating neutrophils fractions (TIN) (q = 0.005, q = 0.0169; **Fig. 5G-H**), consistent with prior observations of subtype-specific immune infiltration patterns ^74–76^. Together, these analyses demonstrate that multiplexed imaging offers a scalable framework for generating spatial insights for research directly from diagnostic samples, bridging routine diagnostics with translational discovery.

## Discussion

Accurate histopathologic diagnosis remains the foundation of precision oncology, particularly in lung cancer, where therapeutic decisions depend on histologic subclassification and biomarker expression. However, current diagnostic workflows are constrained by time-consuming stepwise analyses, evaluating single biomarkers, small sizes of biopsies and the tissue-depleting nature of sequential immunohistochemical (IHC) testing. In this study, we show that multiplexed imaging enables tumor classification and biomarker assessment from a single tissue section, achieving diagnostic performance comparable to conventional workflows while directly addressing these limitations.

A central advantage of multiplexed imaging is its ability to expedite diagnostic analyses by replacing serial single-marker testing with simultaneous detection of dozens of clinically relevant markers. In conventional workflows, histologic subtyping, PD-L1 scoring, and assessment of actionable biomarkers are performed sequentially, causing delays in vital treatment ^77,78^. In contrast, multiplexed imaging enables many of these assessments to be performed in parallel, supporting earlier treatment decision making. In this study, comprehensive evaluation, including diagnosis, PD-L1 scoring, and protein-based assessment of selected actionable targets, was completed within four days; representing a substantial reduction compared to standard clinical timelines ^33,34,36–41^. This consideration is particularly consequential in lung cancer, where most patients present with advanced disease and delays in treatment initiation can adversely affect outcomes ^79^. The panel is flexible and easily accommodates integration of new targets (e.g. MET) as they emerge ^80^.

Multiplexed imaging also conserves limited biopsy material, addressing a persistent challenge in modern diagnostics. Small biopsies are frequently exhausted by serial H&E, IHC, and molecular testing, leading to incomplete diagnosis or biomarker evaluation or repeat invasive procedures. While complementary molecular analysis information may be derived from liquid biopsies, these assays remain costly and further delay treatment initiation. By extracting substantially more information from a single section, multiplexed imaging preserves material for comprehensive molecular profiling, emerging assays, or participation in clinical trials, and reduces the risk of diagnostic failure due to tissue exhaustion. This tissue-sparing property becomes increasingly important as the number of actionable biomarkers continues to expand ^81,82^.

Beyond efficiency and tissue conservation, multiplexed imaging offers conceptual advantages in diagnostic interpretation through complementary mechanisms. First, evaluating all markers on the same tissue section eliminates the need to infer co-expression and spatial relationships across separate sections, a recognized source of variability that is particularly problematic for low-threshold biomarkers such as PD-L1 ^83^. In addition, assessment of multiple lineage and biomarker stains provides contextual information that can resolve diagnostically ambiguous cases. Moreover, multiplexed imaging naturally supports quantitative analysis using automated computational pipelines and facilitates the integration of AI-based tools ^84^. This reduces interobserver variability and enables standardized and reproducible biomarker assessment as shown here for automated PD-L1 scoring in NSCLC and machine-learning–based tumor-infiltrating lymphocytes (TIL) quantification studies ^85–87^.

Importantly, multiplexed imaging also integrates translational and basic research into routine diagnostics without depleting tissue or delaying clinical decision-making. The same assay enables spatially resolved profiling of various markers, generating data for research from every diagnostic sample. It also supports investigation of complex biomarkers that depend on multiple co-expressed markers, such as TILs ^88,89^. This integration of diagnostics and translational research is particularly relevant as neoadjuvant therapies increasingly limit availability of untreated tumor tissue for research.

Several limitations warrant consideration. For some biomarkers, multiplexed imaging provides protein-level surrogates for genomic alterations, and confirmatory testing by NGS remains necessary for definitive genomic characterization in many clinical contexts ^90,91^. In addition, successful implementation depends on the availability of antibody clones compatible with multiplexed protocols. For example, EGFR staining could not be reliably optimized in this study, underscoring the need for continued antibody validation and reagent development. Finally, while this work provides a proof-of-concept, multiplexed imaging should be further developed to penetrate the clinic. Future work should focus on standardizing multiplexed protocols, validating the assay prospectively across institutions, ensuring compatibility with digital pathology systems, and evaluating cost-effectiveness and scalability in high-throughput clinical environments. Integration with machine-learning–based analysis and regulatory-grade automation will be essential for widespread adoption.

While this work focuses on lung cancer, the principles demonstrated here are likely applicable to other malignancies characterized by limited tissue or complex phenotyping requirements. Pancreatic and endometrial biopsies, for example, frequently yield minimal tissue ^92,93^. The ability to assess multiple co-expressed markers may benefit the diagnosis of lymphomas and other hematologic malignancies in which tumor cells may be relatively scarce and surrounded by an inflammatory infiltrate ^94^. Our results suggest that multiplexed imaging may help overcome these challenges by enabling integrated, single-section evaluation of complex marker combinations.

In summary, multiplexed imaging enables fast, tissue-efficient, and accurate diagnostic evaluation while supporting automation and translational research. By consolidating diagnostic and discovery workflows into a single, scalable platform, multiplexed imaging represents a compelling foundation for next-generation pathology in lung cancer and other malignancies characterized by limited tissue and increasing biomarker complexity.

## Materials and methods

### Experimental model and subject details

A cohort of 109 FFPE archival tissues from patients diagnosed with primary or metastatic lung cancer was obtained from Hadassah Hospital Pathology Department. For the calibration cohort, 55 wedge or lobe resection specimens were reviewed by a board-certified pathologist and representative 2 mm cores were compiled into two tissue microarrays (TMAs). For five of the patients, two cores were included in the TMA (#24, #34, #39, #61, #66), and for one of the patients three cores were included in the TMA (#70), resulting in a total of 62 cores. H&E staining was performed on all cores. For the biopsy validation cohort, 54 biopsies were chosen to include a variety of lung tumors, with varying mutations and PD-L1 scores. H&E-stained slides were obtained from Hadassah Hospital Pathology Department archive.

### Data collection

Pathological diagnosis, PD-L1 score, and mutation status were obtained from the medical records.

### Staining and imaging

Tissue sections (5 μm thick) were cut from FFPE tissue blocks of the TMA or biopsies using a microtome, and mounted on Superfrost Plus glass slides (Epredia, Netherlands) for staining. Tissue sections were baked at 70°C for 20 min. Tissue sections were deparaffinized with 3 washes of fresh xylene. Tissue sections were then rehydrated with successive washes of ethanol 100% (2x), 95% (2x), 80% (1x), 70% (1x), and DDW (3x). Washes were performed using a Leica ST4020 Linear Stainer (Leica Biosystems, Wetzlar, Germany) programmed to 3 dips per wash for 30 seconds each. The sections were then immersed in epitope retrieval buffer (Target Retrieval Solution, pH 9, DAKO Agilent, Santa Clara, CA) and incubated at 97°C for 40 min and cooled down to 65°C using Lab Vision PT module (Thermo Fisher Scientific, Waltham, MA). Slides were washed for 5 min with a wash buffer made with DDW, 5% of 20X Stock PBS IHC Wash Buffer with Tween 20 (Cell Marque, Rocklin, CA) and 0.1% (w/v) BSA (Thermo Fisher Scientific, Waltham, MA). Tissue sections were stained and imaged using the COMET (Lunaphore, Switzerland) platform according to the manufacturer’s recommended protocol. Details on the staining conditions for the primary and secondary antibodies are provided in Supplementary Table 1. Multiplexed images underwent pre-processing within the COMET and subsequently in the Horizon software, as implemented by the manufacturer. Images were registered over imaging cycles and background autofluorescence was removed by subtracting an image of a blank channel. For the TMA, multiplexed image sets were extracted by manually marking individual cores using QuPath’s TMA De-arrayer with a diameter of 2.25 mm and a custom Groovy script, which exports them as TIFF files ^95^.

### Diagnosis analysis

Image analysis was performed using QuPath ^95^. Paired H&E and multiplexed images derived from the COMET platform were examined in random order by a board-certified pathologist, who was blinded to the original pathological diagnosis during this examination. The diagnoses given by the pathologist were recorded and compared to the original diagnoses. The degree of agreement between the diagnoses obtained using multiplexed imaging and the original diagnoses obtained using conventional IHC was quantified with Cohen’s kappa coefficient.

### PD-L1 scoring

PD-L1 scores were assigned to each biopsy based on the multiplexed images by a board-certified pathologist and defined as the percentage of viable tumor cells showing membranous PD-L1 staining. PD-L1 scores range between negative (<1% positive cells), weakly positive (1-49% positive cells), and strongly positive (≥50% positive cells) ^96^. First, tumor regions were annotated in each biopsy by a pathologist. For the manual PD-L1 scoring, cell segmentation was performed using QuPath’s cell detection function, using DAPI as the detection channel. Next, PD-L1-positive tumor cells were manually counted, and scores were calculated as the fraction of positive cells out of all segmented tumor cells. For the automated PD-L1 scoring, cell segmentation was performed using InstanSeg ^54^ using DAPI for the nuclear staining and PD-L1, CD15, TIGIT, CD20, Calretinin, Synaptophysin, CD31, CD3, CD30, CD8, CD45, HLA-I, IDO-1, MelanA, CD68, Collagen, CK5, CK20, aSMA, PanCK, Podoplanin, and CK7 as membrane markers. Cell type classifiers were created for each tumor type, using QuPath’s object classifier, to identify the tumor cells based on a threshold depending on the specific tumor marker. Another classifier was created for PD-L1, using a single threshold for all biopsies. Next, for each tumor type, a composite classifier was created by applying both the tumor and PD-L1 classifiers, resulting in an automated quantification of PD-L1-positive tumor cells. Details on the classifier thresholds are provided in Supplementary Table 1.

### TROP2 NMR analysis

Cell segmentation was performed with InstanSeg ^54^ as described above. Tumor cells were classified using the threshold described above. For each tumor cell, TROP2 mean membrane intensity was calculated as the mean signal ±1µm from the cell segmentation. TROP2 mean cytoplasm intensity was calculated as the mean signal within the eroded cell segmentation (shrunk border -1 µm). NMR score was calculated as 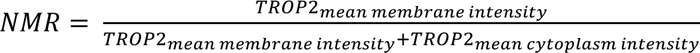. NMR threshold was defined as 0.6. TROP2_NMR+_ samples were defined as samples in which %𝑇𝑢𝑚𝑜𝑟*_NMR_*_≤0.6_ ≥ 75%.

### Microenvironment analysis

Cell segmentation was performed with InstanSeg^54^ as described above. This resulted in 1,734,017 total cells. Tumor and vasculature region masks were generated using the pixel classification module in ilastik^97^. Cells were classified into 14 cell types using CellTune^98^. Cell types and their primary marker definitions are detailed in Supplementary Table 1. Tumor-infiltrating cells were calculated based on the number of cells inside the tumor region masks.

### Ethics

All experiments were conducted in accordance with the Declaration of Helsinki and International Ethical Guidelines for Biomedical Research Involving Human Subjects. The use of the human samples was approved by the local ethics committee of Hadassah Medical Center (0250-23-HMO).

## Acknowledgements

L.K. holds the Fred and Andrea Fallek President’s Development Chair and is supported by the Enoch foundation research fund, the Abisch-Frenkel foundation, the Rising Tide foundation, the Sharon Levine Foundation, Fundación Alberto Palatchi, Dwek center for cancer immunotherapy, and grants from the European Research Council (948811), Israel Science Foundation (2481/20), the Rosetrees Foundation (10004), the Israel Precision Medicine Partnership Program (3830/21), and the Melanoma Research Alliance Team Science Award (1200724). Y.B. is supported by the Clore Scholars Program of the Clore Israel Foundation. E.P. is supported by the Dr. Miriam and Sheldon G. Adelson Medical Research Foundation (AMRF) and the Israel Science Foundation (ISF) Precision Medicine program (1696/20). This study was supported by a grant from the Joint Research Fund of the Hebrew University of Jerusalem and Hadassah Medical Center.

