## Supplementary figures for "Single-section multiplexed imaging enables comprehensive lung cancer diagnosis"

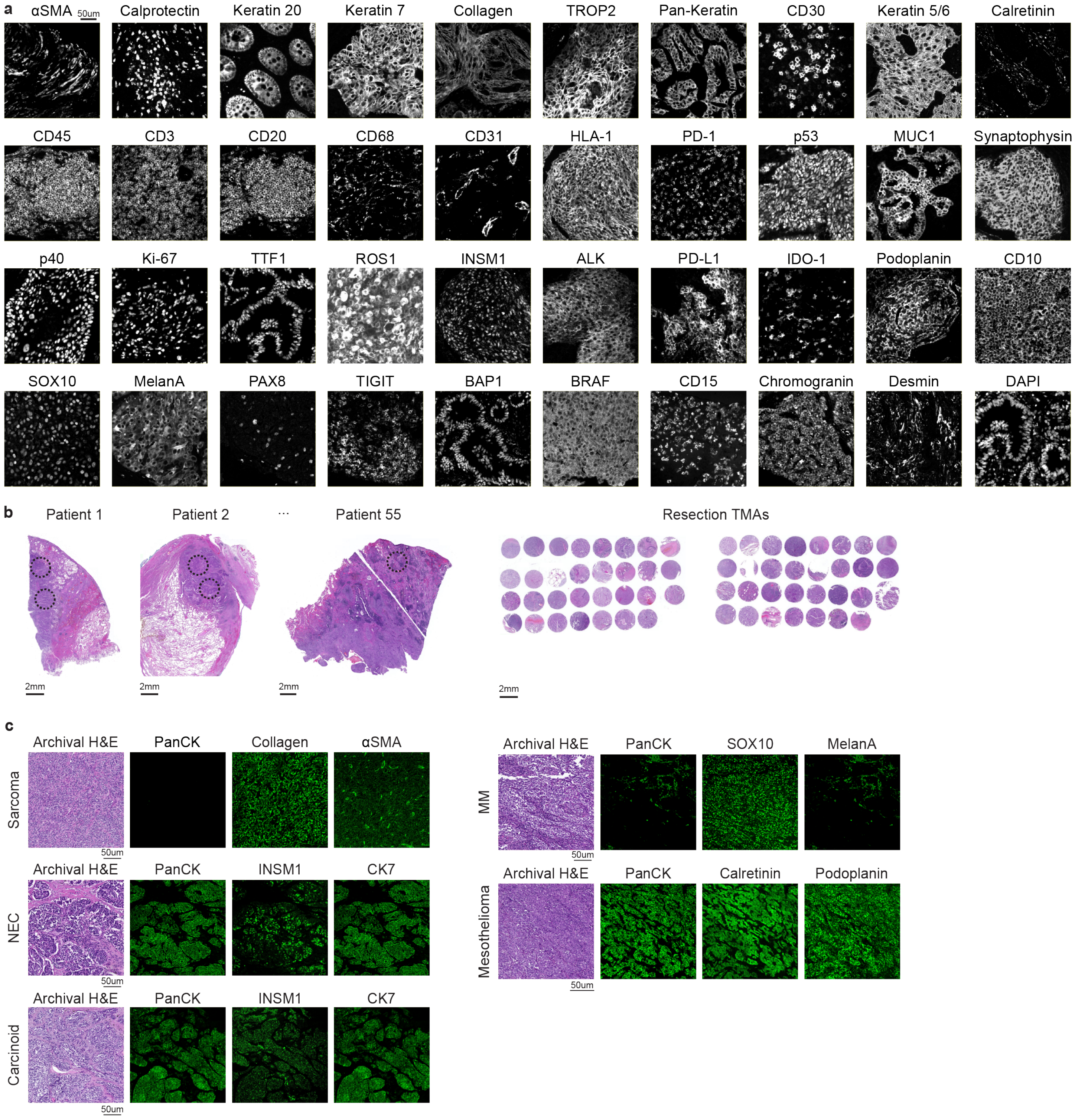


**Supplementary Figure 1: Multiplexed panel calibration on resection TMA**

**(a)** Fluorescent images of samples stained with the panel in Fig. 1B. **(b)** Example H&E images of resection samples (top) marked with areas for core extraction (black circles) used for the generation of a resection TMA (bottom). **(c)** Representative cases demonstrating multiple immunofluorescent stains in addition to standard H&E, leading to a final diagnosis. Abbreviations: NEC = Neuroendocrine carcinoma, MM = Malignant melanoma.

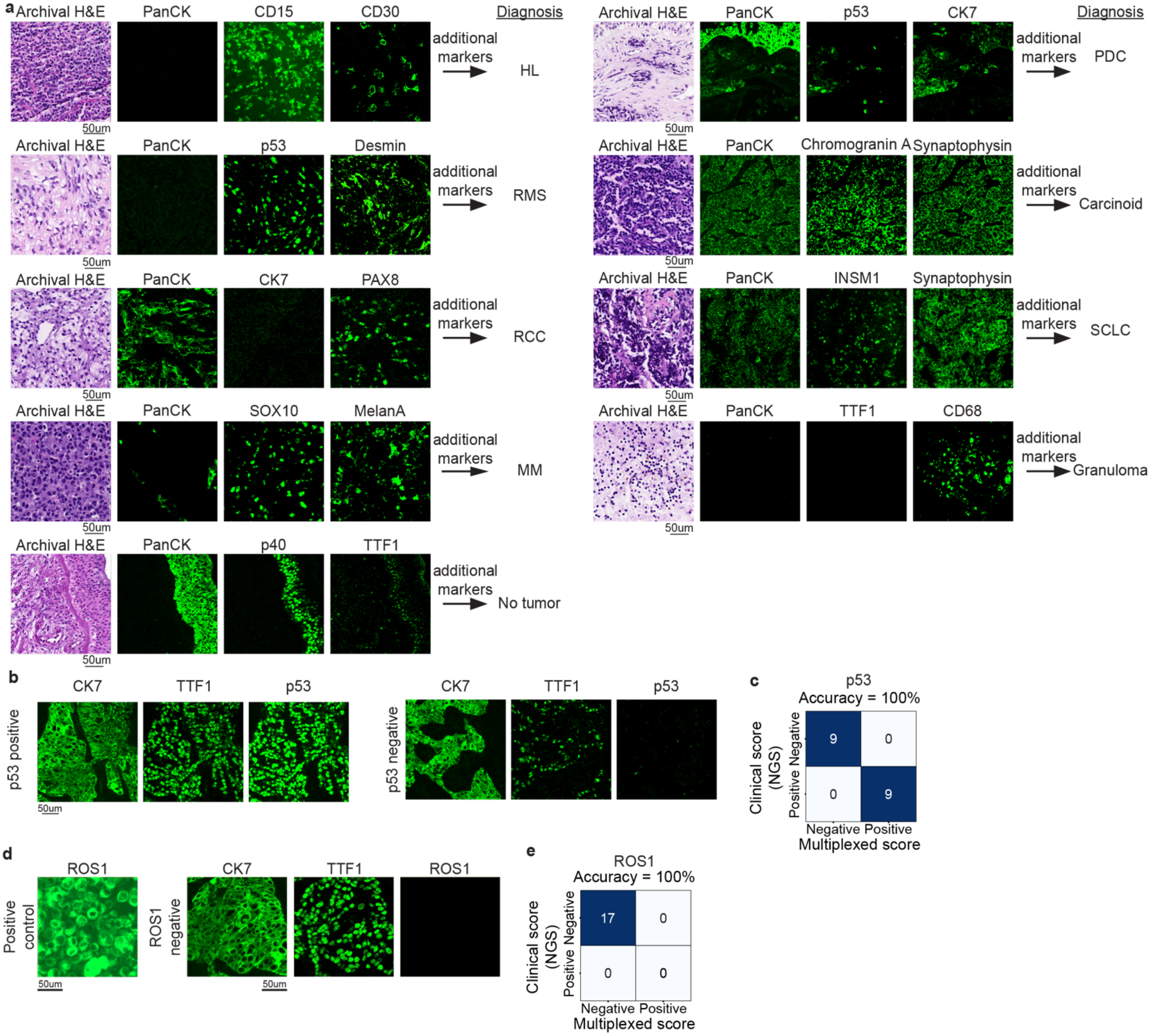


### Supplementary Figure 2

**(a)** Representative cases demonstrating multiple immunofluorescent stains in addition to standard H&E, leading to a final diagnosis. **(b)** Representative images of p53 positive (left) and negative (right) tumors. **(c)** Confusion between NGS (y-axis) and multiplexed image (x-axis) in detecting p53 overexpression. **(d)** Representative images of ROS positive control (left) and ROS negative tumor (right). **(e)** Confusion between NGS (y-axis) and multiplexed imaging (x-axis) in detecting ROS mutation. Abbreviations: HL = Hodgkin lymphoma, RMS = Rhabdomyosarcoma, RCC = Renal cell carcinoma, MM = Melanoma, PDC = Poorly differentiated carcinoma, SCLC = Small cell lung carcinoma.
